# Dynamic evolution of cervical cancer mutations during chemoradiation using novel sampling approach

**DOI:** 10.1101/850388

**Authors:** Bhavana V. Chapman, Tatiana Karpinets, Travis T. Sims, Greyson Biegert, Xiaogang Wu, Andrea Y. Delgado Medrano, Patricia J. Eifel, Anuja Jhingran, Lilie L. Lin, Lois M. Ramondetta, Andrew M. Futreal, Amir A. Jazaeri, Michael Frumovitz, Kathleen M. Schmeler, Jingyan Yue, Aparna Mitra, Kyoko Yoshida-Court, Travis Solley, Geena Mathew, Mustapha Ahmed-Kaddar, Jianhua Zhang, Ann H. Klopp, Lauren E. Colbert

## Abstract

**Objective:** The aim of this study was to validate a whole exome sequencing approach to longitudinally characterize the tumor mutational profile of cervical cancer patients undergoing chemoradiation (CRT).

**Experimental Design:** Cervical cancer tumor specimens from twenty-seven patients undergoing chemoradiation were collected before and throughout CRT and whole exome sequencing (WES) was performed to characterize individual mutations and alterations in unique genes. WES data were analyzed from cervical cancer patients in The Cancer Genome Atlas (TCGA) as a comparison group.

**Results:** Over 93% of mutated genes detected at baseline were present in TCGA. Tumor purity from collected swabs correlated with MRI tumor volumes during the course of treatment (R^2^=0.969). CDK4/CDK6/cyclin D1-related gene mutations involved in the ERK1/2, p16INK4, and p53 pathway and G1/S checkpoint most commonly persisted at the end of CRT.

**Conclusion:** This non-invasive swab technique to serially sample tumor during CRT will allow new discoveries of dynamic tumor mutational profile changes during chemoradiation for mucosal tumors. Mutations that survived or increased during the initial weeks of radiation treatment are potential drivers of radiation resistance including the CDL4/CDK6/cyclin D1-related pathway.

**Statement of Translational Relevance:** There are no established biomarkers to predict chemoradiation (CRT) response for cervical cancer patients. Serial biopsies cannot be performed due to risks of bleeding and fistula. We used a novel non-invasive swab-based biopsy technique to obtain serial samples from a cohort of twenty-seven patients through the course of treatment, and validated this approach to obtain whole exome sequencing data. We analyzed dynamic tumor mutation profiles during CRT. Results from this study show that mutations in CDK4/CDK6/cyclin D1-related genes increased at the end of CRT, suggesting this pathway as a potential driver of radiation resistance.

## Introduction

Cervical cancer continues to be one of the most common gynecologic cancers among women globally.^1^ Approximately 13,000 new cases of invasive cervical cancer will be diagnosed in the United States in 2019, resulting in more than 4,000 deaths.^2^ It is well established that persistent exposure to high-risk human papilloma viruses (HPVs) plays an essential role in cervical dysplasia and carcinogenesis, causing the vast majority of cervical cancer.^3^ The global burden of cervical cancer is growing despite the development of HPV vaccines aimed at preventing the disease. Vaccines currently have a low adoption rate, and it will undoubtedly be decades before cervical cancer rates are significantly reduced.^4^ Consequently, multimodality therapy with chemoradiation (CRT) continues to be the standard of care for treating locally advanced disease.^5^ As a result, novel therapeutic targets directed at cervical cancer could expand upon current strategies to combat this gynecologic malignancy.

Past studies have identified somatic mutations in *PIK3CA, PTEN, TP53*, and KRAS in cervical carcinoma.^6–8^ But in recent years, next generation sequencing has notably increased our understanding of cervical cancer biology.^9,10^ Whole exome sequencing (WES) on cervical cancer samples provides complementary, pathway-generating insight on somatic mutations, mutational burden, and single nucleotide human exome variants (SNVs) that may be functionally significant.^9^ Several studies have evaluated these genomic alterations in cervical cancer. In cervical squamous cell carcinoma, somatic mutations include recurrent *E322K* substitutions in the *MAPK1* gene, inactivating mutations in the HLA-B gene, and mutations in *EP300, FBXW7, NFE2L2, TP53* and *ERBB2*.^9^ The Cancer Genome Atlas (TCGA) project identified *SHKBP1, ERBB3, CASP8, HLA-A* and *TGFBR2* as novel significantly mutated genes.^10^ Another study used WES to identify 64 somatic mutation genes in 3 cervical tumors and found that the HPV16-positive tumors had fewer somatic mutated genes than HPV-negative tumors, which was validated in the TCGA dataset.^11^ In cervical small cell neuroendocrine tumors, *ATRX, ERBB4*, and genes in the Akt/mTOR pathway were most frequently mutated on WES.^12^ Though recent studies have recognized mutation prevalence and a variety of gene expression signatures, none of the studies to date have conducted WES on longitudinal tumor samples over the course of treatment. It remains unclear what mutational differences may predict and distinguish treatment response.

Here we report serial whole exome sequencing of 27 locally advanced cervical cancer tumors. Our goal was to longitudinally characterize the mutational profile of cervical cancer samples from a cohort of patients undergoing CRT. We hypothesized that mutations present at baseline may change in real time throughout the course of CRT. Previously, serial biopsies during CRT have been impossible to obtain due to risks of bleeding and fistula, but our novel non-invasive swab-based biopsy technique allows us to serially sample and analyze the tumor mutational profile during the course of treatment. The goal of this study was to validate the use of this non-invasive technique to demonstrate serial changes in mutations, and, in turn, identify potential mutations associated with therapeutic resistance.

## Materials and Methods

### Patient population and treatment characteristics

Patients were enrolled in an IRB approved (2014-0543) multi-institutional prospective clinical trial at The University of Texas MD Anderson Cancer Center and the Harris Health System, Lyndon B. Johnson General Hospital Oncology Clinic (**Figure 1A**). This study was designed to evaluate metagenomic changes in the cervical and intestinal microbiota during chemoradiation. Inclusion criteria were newly diagnosed locally advanced cervical cancer per the Federation of Gynecology and Obstetrics (FIGO) 2009 staging system. Patients with clinical stage IB1-IVA, visible, exophytic tumor on speculum examination with planned treatment of intact cervical cancer with definitive radiation therapy, including external beam radiation therapy and brachytherapy with concurrent cisplatin were included. Patients with any previous pelvic radiation therapy or treatment for cervical cancer, including transvaginal cone irradiation, were excluded.

**Figure 1.**
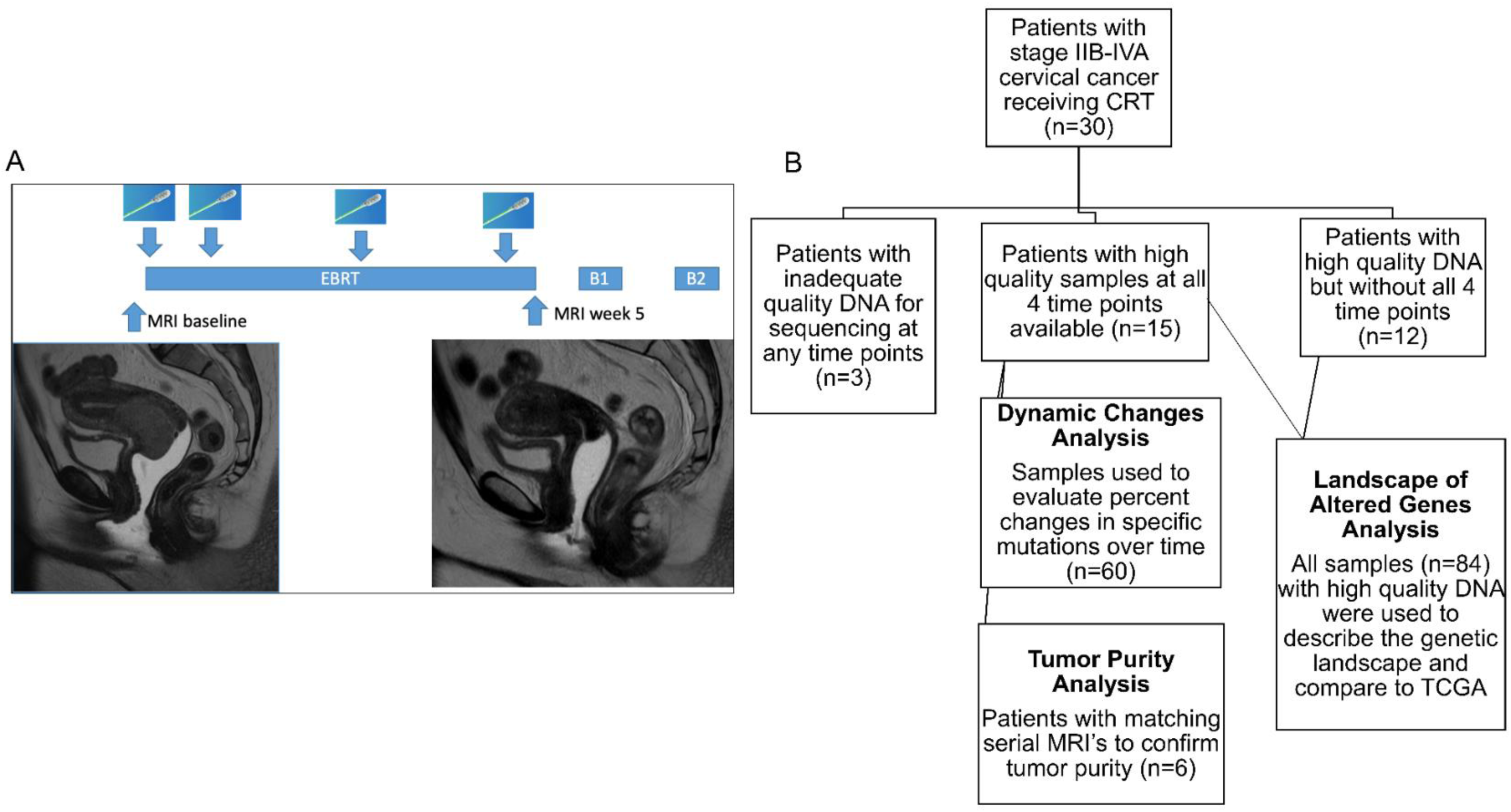
Overall study schema for patient sample collection and analysis. (A) Patients with locally advanced cervical cancer undergo 5 weeks of external beam radiation therapy (EBRT) followed by two brachytherapy treatments (B1 and B2), with samples collected at Baseline, Week 1, Week 3 and Week 5 of radiation therapy. Patients also underwent optional MRI imaging at baseline and at week 5. (B) 30 patients were accrued on protocol, and of these, 27 patients with 85 total samples had adequate DNA quantity and quality for sequencing. These samples were used to describe the overall mutational landscape of the population. 15 patients had samples available at all 4 timepoints, and these samples were used to analyze dynamic changes over time. 6 patients had matching samples at all timepoints and serial MRI’s, and these samples were used to validate the sequencing technique using tumor purity analysis.

Patients underwent the standard-of-care pretreatment evaluation for their disease, including a tumor biopsy to confirm diagnosis; pelvic magnetic resonance imaging (MRI) and positron emission tomography/computed tomography (PET/CT); and standard laboratory evaluations, including a complete blood cell count, measurement of electrolytes, and evaluation of renal and liver function. HPV-positivity was confirmed by p16 staining of tumor biopsies on initial institutional pathologic review. Patients received pelvic radiation therapy to a total dose of 40 to 45 Gy delivered in daily fractions of 1.8 to 2 Gy over 4 to 5 weeks. Following completion of pelvic radiation therapy, patients received intracavitary brachytherapy with pulsed-dose or high-dose rate treatments over 44 to 48 hours or the equivalent with high-dose-rate brachytherapy. Patients received cisplatin 40 mg/m^2^ weekly delivered according to standard institutional protocol. Patients underwent repeat MRI at the completion of external beam radiation therapy or at the time of brachytherapy, as indicated by the extent of disease.

### Sample collection and DNA extraction

Isohelix swabs (product # DSK-50 and XME-50, www.isohelix.com, UKSamples) were brushed against the viable cervical tumor several times by an attending radiation oncologist or gynecologic oncologist at either The University of Texas MD Anderson Cancer Center or Harris Health, Lyndon B. Johnson Hospital. The isohelix swab has a unique matrix design that permits the yield of one to five mcg of high quality DNA sufficient for sequencing applications from a single swab of the tumor surface^13^. Patients underwent swabs at baseline, the end of week 1 (after 5 fractions), at the end of week 3 (after 10-15 fractions), and within a week prior to the first brachytherapy treatment or at the time of brachytherapy (week 5), for a total of four swabs during radiation therapy. Attention was taken to serially swab the same general tumor region. DNA was extracted from normal buccal and cervical cancer samples per Isohelix # DSK-50 manufacturer’s instructions.

### Whole exome sequencing and mutational analysis

The Illumina WES assay was optimized for normal buccal control and cervical tumor DNA swab samples. Captured libraries were sequenced on a HiSeq 2000 series (Illumina Inc., San Diego, CA, USA) on a TruSeq v3 Paired-end Flowcell according to manufacturer’s instructions at a cluster density between 700–1000K clusters/mm2. Sequencing was performed on a HiSeq 2000 series for 2 × 100 paired end reads with a 7 nucleotide read for indexes using Cycle Sequencing v3 reagents (Illumina). All regions were covered by >200 reads.

WES exome sequencing data were aligned to human reference genome using BWA.^14^ Duplicate reads were removed using Picard and then realigned and recalibrated by the Genome Analysis Toolkit.^15^ Samples used for each analysis are outlined in **Figure 1B**. Tumor purity was calculated from single nucleotide variants by TPES.^16^ MuTect and Pindel were applied to detect somatic SNVs and small InDels using buccal DNA samples from each patient as normal controls.^17,18^ Variants were classified into 3 categories: somatic, germline, and loss of heterozygosity based on variant allele frequencies in the tumor and matched normal buccal samples. Somatic copy number alterations reporting gain or loss of each exon were noted using a previously described algorithm.^19^ COSMIC, TCGA, and dbSNP databases were used to identify potential functional consequences of detected variants which were annotated using VEP, Annovar, CanDrA.^20–23^

Cervical tumors were contoured on serial MR imaging for six patients during the course of CRT using RayStation. 3-dimensional volumes were generated from contours. Regression analysis was utilized to evaluate the goodness of fit between tumor purity (previously described) versus MRI tumor volumes.

### Comparison against published databases

TCGA mutations were downloaded using the TCGABiolinks package in R. The mutect and indel pipeline output of the most recent version of the “CESC” dataset was used. The NCI Genomic Data Commons (GDC) Data Portal was also used to query submitted cervical cancer samples with available mutation data.

### Pathway enrichment analysis

To query for potential radiation-related genes, the list of altered genes in all patient samples was queried using the Clarivate Analytics MetaCore database.

### Selection of potential drivers of radiation resistance

The mutations and specific genes that had the greatest percent change from baseline in an individual patient were summed to obtain an average change in the population. These mutations and genes were selected as potential drivers of radiation resistance and pathway analysis was performed using MetaCore database.

### Data Sharing

All code used for these analyses and de-identified raw datasets of mutations will be shared at github.com/ColbertLab/Whole-Exome-Seq-Paper.

## Results

### Patient and tumor characteristics

We performed WES on cervical samples from 27 cervical cancer patients. Tumor samples passing quality control were collected from 15/27 patients at all four sampling timepoints. Clinico-pathologic data are summarized in **Table 1**. Overall, approximately 63% of the patients (17/27) had advanced stage disease (stage IIB or greater) and the majority of patients had squamous cell carcinoma with moderate or poor differentiation. With respect to HPV status, HPV16 was the most frequent genotype (40.7%), followed by HPV18 (7.4%), and other high-risk HPV strains (25.9%). HPV status was unknown in 25.9% of study patients. On pre-treatment MRI, the largest tumor dimension and highest node level were identified. Using the short axis diameter, we found that the median cervical tumor size was 4.6 cm (range 2.2-11.5 cm). Nineteen patients (70%) were identified as having positive pelvic or para-aortic lymph nodes by positron emission tomography (PET) or computerized tomography (CT) scan.

### Baseline somatic mutational landscape

The distribution of detected mutations for each patient across the 4 timepoints is shown in **Figure 2A**, while detected mutations for each patient with respect to each timepoint is shown in **Figure 2B**. The majority of baseline mutations were nonsynonymous. In evaluating exonic mutations across timepoints, nonsynonyous mutations were more prevalent in tumor samples whereas stopgain and stoploss mutations were less abundant **(Figure 2C)**. Breakpoints of large deletions and medium-sized insertions from paired-end short reads were used as a means to detect frameshift/nonframeshift and insertions/deletions **(Figure 2D)**.

**Figure 2.**
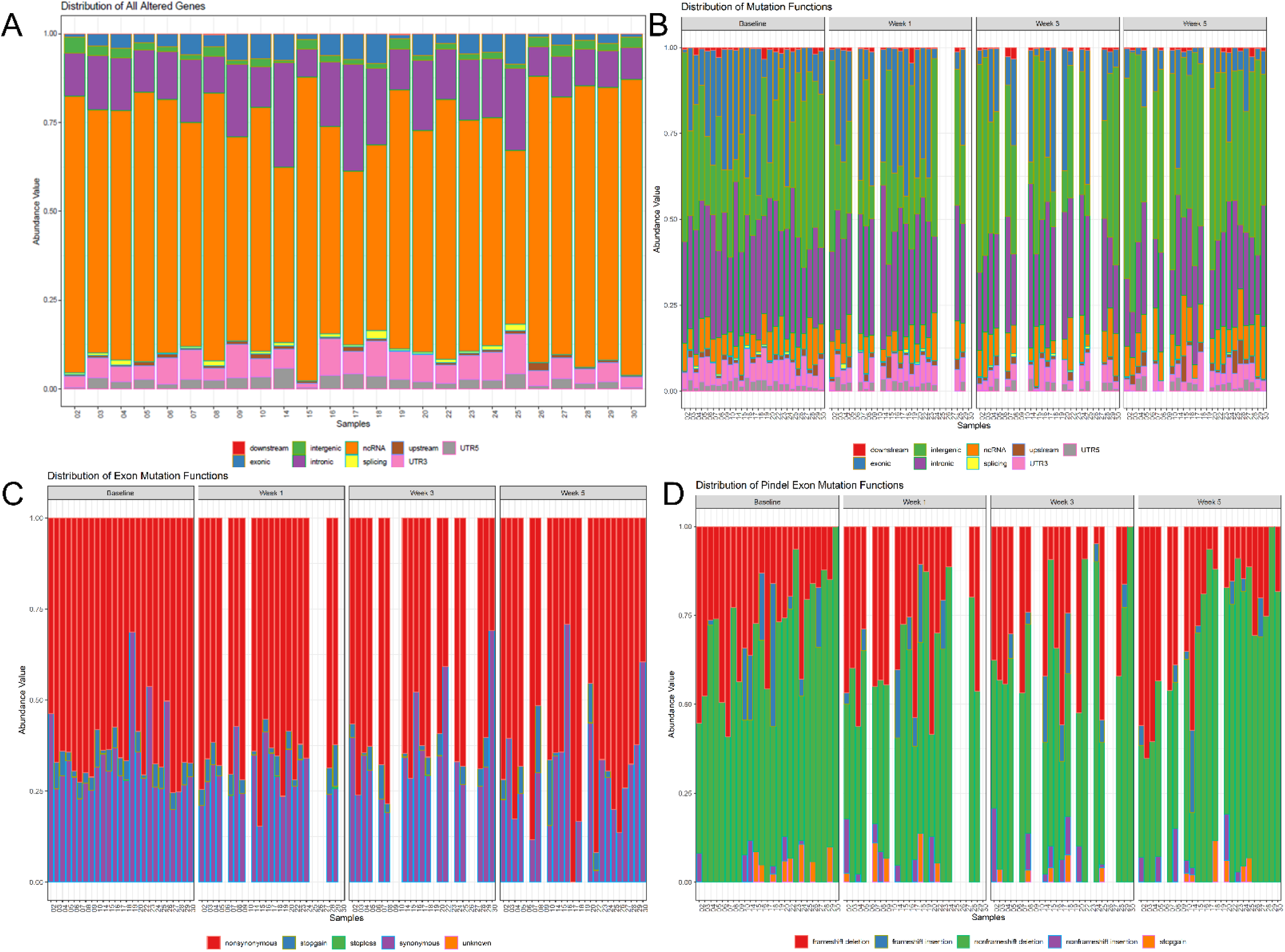
Baseline mutation type stacked bar plots (TCGA vs our population). (A) Stacked bar chart displaying the relative abundance of mutation events in all genes for each patient across all available samples throughout treatment. Similar to (A), (B) highlights the relative abundance of mutation events in all genes for each patient at separate timepoints during treatment. Thus showing that exon mutations as critical contributors to the mutation burden. Figure (C) then displays the relative distribution of exon specific mutation functions for each patient, at separate timepoints during treatment. Additionally, figure (D) shows the relative abundance of insertion/deletion mutations present in exons for each patient stratified by time point. For figures B-D, patient samples with missing data are represented by empty bars.

All substitutions, insertions and deletions were filtered for exonic, nonsynonymous stop/gain and stop/loss alterations. The entire list of mutations is available in **supplemental table 1**. This overall landscape was very similar to known TCGA mutations in the most recent version of the CESC dataset **(Figure 3)**. Overall, 94% (1339/1430) detected exonic, nonsynonymous, stop/gain or stop/loss gene alterations in our samples were present in the TCGA CESC dataset **(Figure 3A). Supplemental table 2** lists all overlapping altered genes between TCGA CESC and our samples overall and at each individual timepoint. **Supplemental table 3** and **supplementary table 4** list the frequencies of all gene alterations in TCGA **(Figure 3B)** and our sample dataset **(Figure 3C)**, ranked by frequency. Among the top 30 most frequently mutated genes, changes in *TTN, MUC4, KMT2C, LRP1N, PIK3CA*, and *KMT2C* and *KMT2D* were present in both groups. There were some unique alterations to our population, including *CDC27*, an important gene for radiation response. Baseline pathway enrichment analysis using ranking of the most common mutations indicates that mutations involved in the PI3K/Akt and Wnt/beta-catenin pathways were commonly detected, consistent with known cervical cancer mutations (**Figure 3D; supplementary table 5)**.

**Figure 3.**
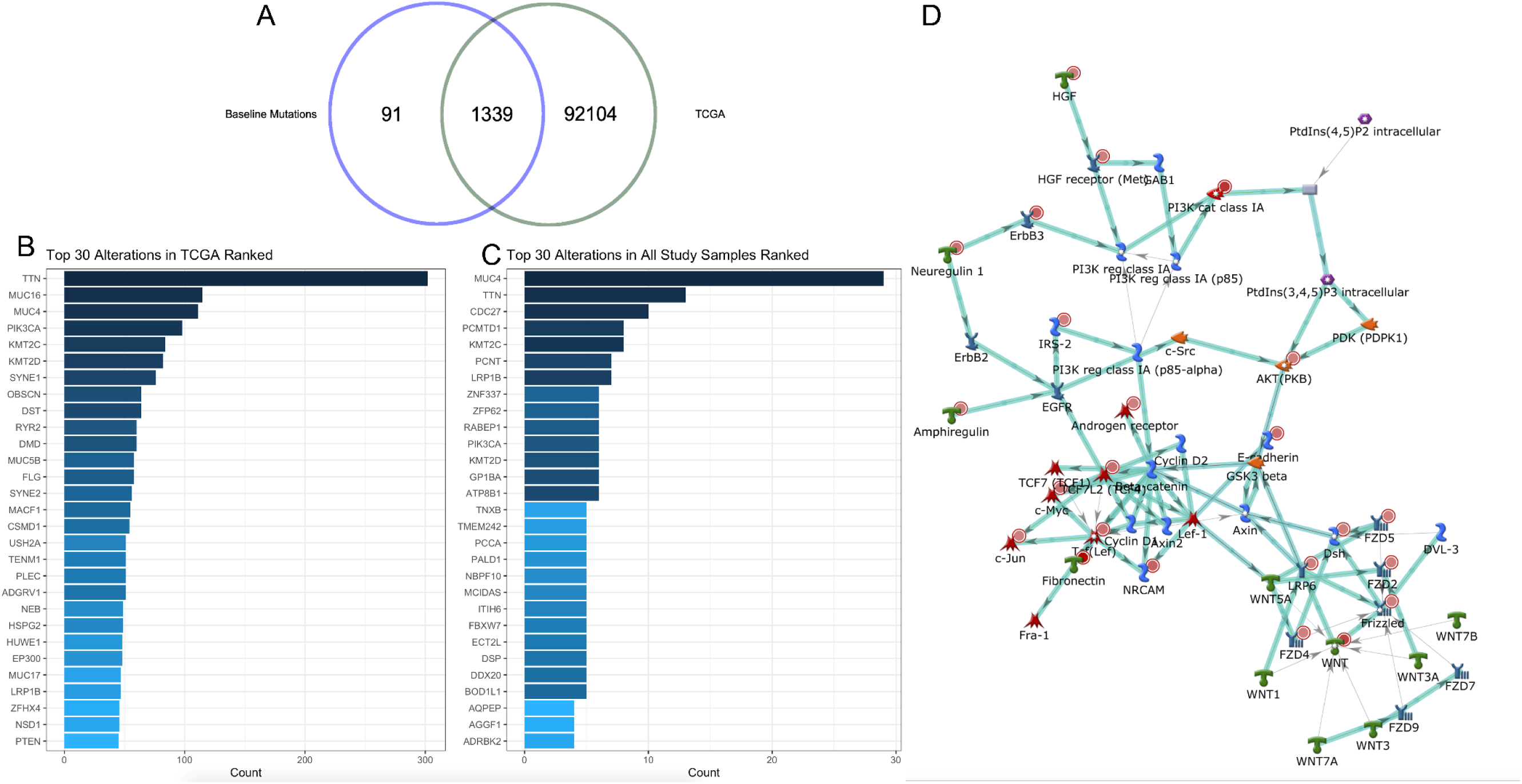
Overall Gene Alterations from All Samples is similar to landscape of TCGA cervical squamous cell carcinoma dataset. (A) 94% of (1339/1430) altered genes in baseline samples (defined as substitutions, insertions or deletions in gene), were also identified in the TCGA dataset, suggesting accurate identification of cervical cancer related genes. (B) Distribution of top 30 most altered genes in our study sample and TCGA (C) is also similar. (D) Pathway analysis was performed of top 100 altered genes in the study samples at baseline, and the most frequency pathway is shown here, centered on PI3K and related genes, again consistent with expected mutational landscape for cervical cancer.

### Evolution of tumor mutational changes

For patients who had adequate DNA at all timepoints, baseline exonic, non-synonymous, stop/gain and stop/ loss mutations were filtered, and changes in frequency of these alterations over time was plotted **(Figure 4A)**, demonstrating that overall, mutations present at baseline decrease over time, although some persist. Ranking of altered genes changed from baseline (**Figure 4B**) to week1 (**4C**), week 3 (**4D**) and week 5 (**4E**), suggesting that there is dynamic evolution throughout chemoradiation. As a whole, mutations in *MUC4* persisted throughout the course of chemoradiation.

**Figure 4.**
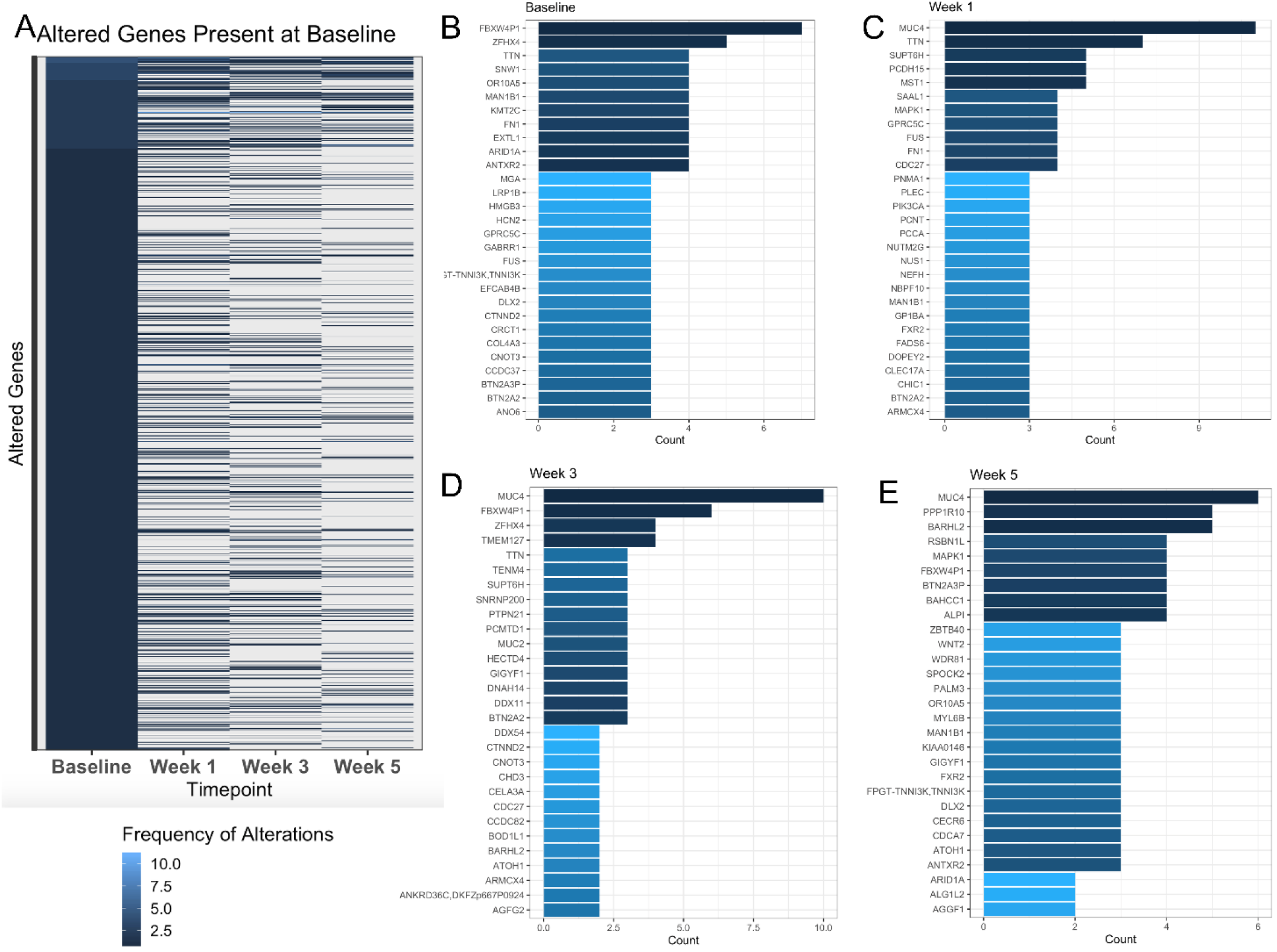
Changes in gene alterations over time for patients with all 4 time points (N=15) demonstrated overall decrease over time, but shifts in specific alterations. (A) All altered genes (exonic non-synonymous mutations, including substitutions, insertions and deletions) present at baseline decrease throughout the course of CRT. Alterations present at Baseline (B), Week 1 (C), Week3 (D), and Week 5 (E)

### Changes in MRI tumor volumes and mutational profile in a patient with persistent tumor

Given this non-invasive biopsy technique, in order to quantify changes in allele frequency for low quality DNA, we compared calculated tumor purities to known tumor volume at each time point. **Figure 5A** demonstrates serial MRI images taken at baseline and week 5 for one poor responder, with tumor contoured in gray. 3-dimensional tumor volumes were calculated at each timepoint for the 6 patients with serial MRI’s. There was a high degree of linear correlation (**Figure 5B**, R^2^=0.969) between computed tumor purity and known tumor volume. Once this was confirmed, calculated percent change in allele frequencies of individual mutations over time were calculated (**supplemental table 6**). All significantly changed mutations in individual patients were then summed and averaged to determine changes in individual mutations for the whole population **(Figure 5C; supplemental table 7).** The top 20 ranked genes by change at week 5 included *TCF25, MUC2, PARP10* and *PIK3CA* **(Figure 5D)**. Pathway analysis of the ranked mutated genes present at week 5 demonstrate perturbations in the G1/S transition via CDK4/CDK6/cyclin D1-related gene mutations, a key radiation sensitivity pathway (**Figure 5E**).

**Figure 5.**
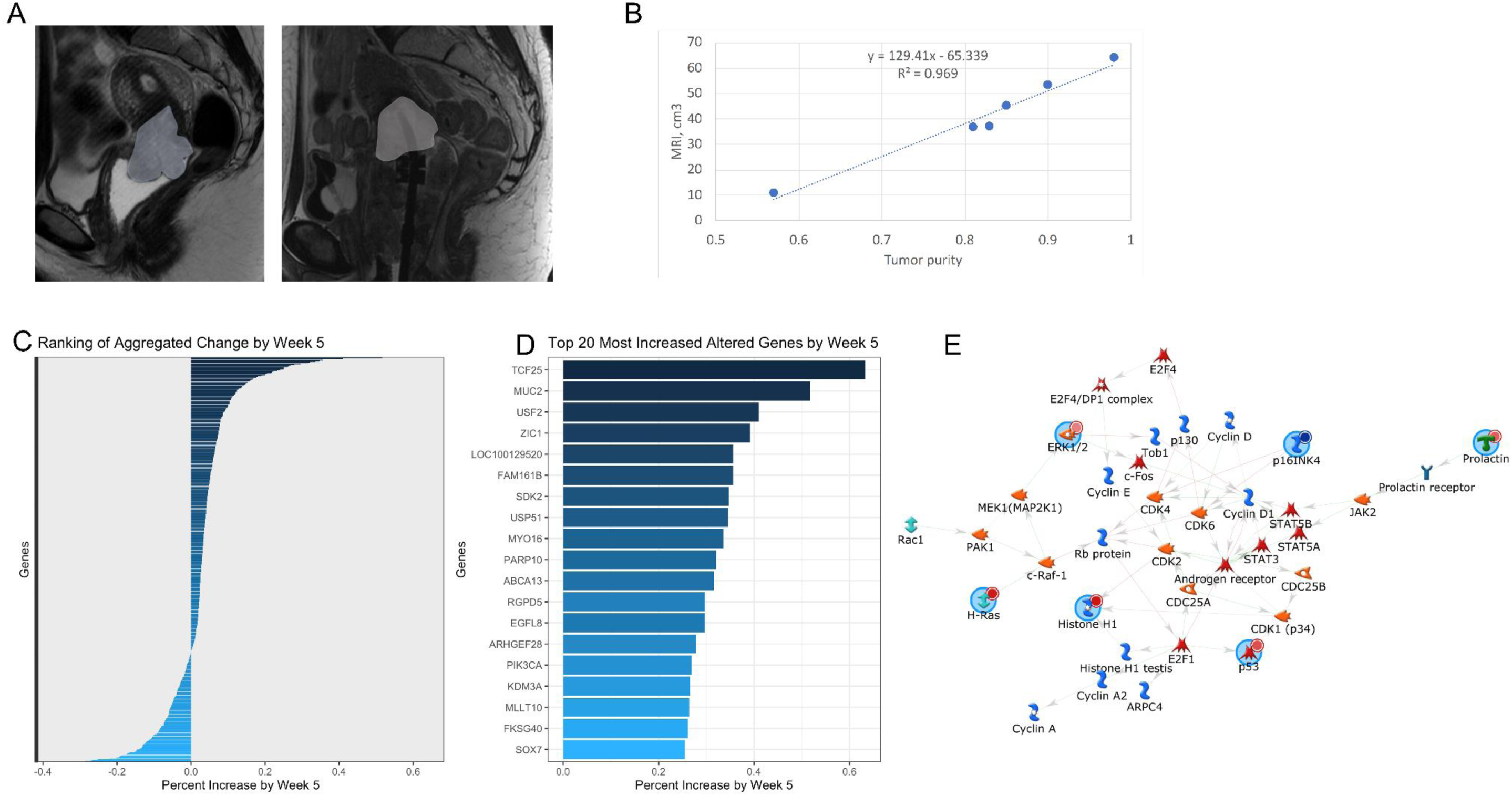
Individual mutations, insertions and deletions are highly dynamic throughout the course of CRT. (A) Demonstrates a representative poor responder with volume contoured to obtain change in overall tumor volume. (B) Demonstrates linear correlation between calculated tumor purity and MRI volume, used to adjust for relative mutation burden. (C) Ranked percent change in allele frequencies for individual genes summed by gene, demonstrating both increased and decreased allele frequencies from baseline to week 5. (D) Top 20 most increased altered genes by week5 in the overall population, suggesting potential drivers of radiation resistance. (E) The highest ranked pathway, according to increase over time and summed number of patients, centered on CDK4/CDK6/ and p16INK4 with included nodes highlighted in blue.

## Discussion

To our knowledge, this is the first study to serially sample cervical cancer tumors and comprehensively characterize alterations in the mutational landscape using whole exome sequencing during the course of CRT. Obtaining tumor biopsies during the course of treatment is painful for patients and logistically challenging. Historically, the aforementioned limitations pose significant barriers to evaluating dynamic changes in tumor genetic signatures in response to ongoing treatment. Our novel, non-invasive swab-based biopsy technique which allows for DNA collection from cervical brushings during pelvic exam circumvents these obstacles. We noted that tumor purity correlated with residual cervical tumor volumes measured from MRI performed at corresponding timepoints. Moving forward, this method could be applied to other gynecologic, head and neck, anorectal, and skin malignancies to establish biomarker predictors for treatment response and potential drivers of treatment resistance which may be undetectable from the analysis of initial biopsies only.

The distribution of somatic mutations detected in our patient cohort is similar to other patient cohorts which further validates our findings. There was a high degree of overlap between the mutated genes detected at baseline in our patient samples and the TCGA cervical cancer cohort which is comprised of chemoradiation-naive patients. *MUC4*, the most frequently mutated gene in our cohort, encodes a membrane-bound mucin with promotes tumorigenicity through epithelial invasion, immune surveillance evasion, and suppression of apoptosis.^25^ It is implicated in cervical squamous dysplasia and carcinoma.^26^ MUC4 expression was found to be inversely related to survival in patients with epithelial carcinomas.^27^ *MUC16*, another mucin membrane protein, encodes the extracellular domain of CA125 and is commonly mutated in ovarian and gastric cancer.^32^ *MUC16* is implicated in aberrant cell proliferation via the JAK-STAT pathway and EGFR signaling with activation of downstream Akt and ERK1/2.^33^ *TTN*, another frequently mutated gene detected in our cohort and TCGA, encodes for Titin, the largest protein found in humans encoded by 364 exons.^28^ Mutations in *TTN* have been reported in dilated cardiomyopathy and Titin autoantibodies have been detected in patients with Sjögren’s syndrome and systemic scleroderma.^29,30^ Patients with mutations in *TTN* have been shown to possess higher tumor mutational burden and improved response to immune checkpoint blockade therapy in various solid tumors of skin, lung, stomach compared to those with wild-type *TTN.*^31^

Pathway analysis of baseline mutations in our cohort demonstrates that the MAPK mitogenic pathway was the most commonly perturbed pathway overall. Phosphoinositide 3-kinase (PI3K) pathway-related mutations were also prevalent. Others have reported perturbations in *PIK3CA*, an oncogene part of the ERBB2/PI3K/AKT/mTOR pathway with a battery of druggable targets, present in over 50% of cervical tumor and cervical cancer cell lines, which is consistent with our findings.^3^ Schuurbiers et al. previously described PI3K-Akt pathway activation and mechanisms of radiation resistance in non-small cell lung cancer which could similarly be involved in cervical cancer.^35^ Additionally, *in vitro* studies demonstrate the *PIK3CA* E545K mutation results in resistance to platinum chemotherapy.^36^ The E545K mutation is the single most common mutation among TCGA cervical cancer patients and is present at a rate of 13% (38/289).

When examining mutations that persisted at the end of radiation, those involved in the G1/S transition involving ERK1/2, p16INK4, and p53 were most frequently mutated and driven by several CDK4/CDK6/cyclin D1-related gene mutations. The G1/S checkpoint is involved in the DNA damage repair process and radiation-induced cell death.^37^ The aberrant CDK4/CDK6/cyclin D1 pathway is well known in the pathogenesis of HPV-associated malignancies and CDK4/6 inhibitors are even being explored as potential therapeutic agents in the treatment of HPV-negative cervical cancer.^38,39^

Presently, there is a paucity of data examining serial genomic profiling of tumors before and after treatment, especially in the locally advanced setting. In the TCGA cervical cancer cohort, over 60% (138/228) patients had FIGO stage I disease, and over 75% (106/138) of which were treated with upfront hysterectomy.^10^ The majority of these early stage cervical cancer patients likely did not require adjuvant therapy. Thus, the baseline mutational profile of patients from TCGA may not be directly comparable to patients with locally advanced cervical cancer who go on to receive definitive chemoradiation as described in our cohort. WES feasibility studies have been described in the recurrent and metastatic setting, but studies examining *de novo*, disease site-specific populations are limited.^40^ A recent series of 28 patients with rectal cancer reported WES analysis before and after preoperative CRT demonstrated persistent mutations in *CTDSP2, APC, KRAS, TP53* and *NFKBIZ* conferred treatment resistance.^41^ Next-generation sequencing has been described to identify gene signatures predictive of treatment response from initial biopsies and surgical specimens after preoperative radiotherapy for early stage breast cancer (PMID31525407).

In conclusion, this study provides evidence that a novel swab-based technique can be utilized to serially sample cervical tumors for whole exome sequencing analysis. We posit that mutations that survive the initial weeks of radiation treatment may be clinically relevant drivers of radiation resistance. Future work will focus on characterizing the clonal architecture of residual tumors to identify granular molecular signatures predictive of treatment response.

## Author Contribution

All authors were involved with subject identification and data collection, interpretation of statistical analysis, and review and approval of final manuscript. Study concept was developed by AHK, LEC and BVC. Analyses were performed by LEC, THK, XW, GB. Drafting and editing of manuscript was performed by all authors.

## Acknowledgements

This work was supported in part by the Radiological Society of North America (RSNA) Resident/Fellow Award (LEC), National Institutes of Health (NIH) through MD Anderson’s Cancer Center Support Grant P30CA016672 and the NIH T32 grant #5T32 CA101642-14 (TTS) and The University of Texas MD Anderson Cancer Center HPV-related Cancers Moonshot (LEC, AHK). The human subjects who participated in this study are gratefully acknowledged, in addition to our clinical research team, and scientific publications team.

---

| Characteristic | No. of patients | % |
| --- | --- | --- |
| **Age at diagnosis, years** |  |  |
| Median | 47.5 |  |
| Range | 30-72 |  |
| **Race/Ethnicity** |  |  |
| African American | 2 | 7.4 |
| American Indian/Alaskan Native | 1 | 3.7 |
| Asian | 1 | 3.7 |
| Hispanic/Latino | 16 | 59.3 |
| White/Caucasian | 7 | 25.9 |
| **Histology** |  |  |
| Adenocarcinoma | 8 | 30.0 |
| Squamous cell carcinoma | 19 | 70.0 |
| **2009 FIGO stage** |  |  |
| IB1 | 3 | 11.1 |
| IB2 | 5 | 18.5 |
| IIA | 2 | 7.4 |
| IIB | 12 | 44.4 |
| IIIB | 4 | 14.8 |
| IVA | 1 | 3.7 |
| **Tumor grade** |  |  |
| I | 2 | 7.4 |
| II | 11 | 40.7 |
| III | 9 | 33.3 |
| Unknown | 5 | 18.5 |
| **Highest clinically involved nodal level on PET or CT** |  |  |
| Para-aortic | 4 | 14.8 |
| Common iliac | 3 | 11.1 |
| External iliac | 10 | 37.0 |
| Internal iliac | 2 | 7.4 |
| Node-negative | 8 | 29.6 |
| **Max tumor dimension on MRI, cm** |  |  |
| Median | 4.6 |  |
| Range | 2.2-11.5 |  |
| Unknown | 1 |  |
| **HPV genotype** |  |  |
| HPV16 | 11 | 40.7 |
| HPV18 | 2 | 7.4 |
| Other high-risk HPV | 7 | 25.9 |
| Unknown | 7 | 25.9 |
| **Smoking status** |  |  |
| Current | 3 | 11.1 |
| Former | 12 | 44.4 |
| Never | 12 | 44.4 |

